# Transcriptional Profiling of Patient Isolates Identifies a Novel TOR Regulatory Pathway in Cryptococcal Virulence

**DOI:** 10.1101/368738

**Authors:** Yoon-Dong Park, Joseph N. Jarvis, Guowu Hu, Sarah E. Davis, Jin Qiu, Nannan Zhang, Christopher Hollingsworth, Angela Loyse, Paul J. Gardina, Tibor Valyi-Nagy, Timothy G. Myers, Thomas S. Harrison, Tihana Bicanic, Peter R. Williamson

**Affiliations:** Laboratory of Clinical Immunology and Microbiology, National Institute of Allergy and Infectious Diseases, National Institutes of Health, Bethesda, MD, USA; Botswana-UPenn Partnership, Gaborone, Botswana; Division of Infectious Diseases, Department of Medicine, Perelman School of Medicine, University of Pennsylvania, Philadelphia, PA, USA; Department of Clinical Research, Faculty of Infectious Diseases and Tropical Medicine, London School of Hygiene and Tropical Medicine, London, UK; Global health group, Department of Infection and Immunity, St George’s University of London, UK; Genomic Technologies Section, Research Technologies Branch, National Institute of Allergy and Infectious Diseases, National Institutes of Health, Bethesda, MD, USA; Department of Pathology, University of Illinois at Chicago, Chicago, IL, USA; Institute of Infection and Immunity, St George’s University of London, London, UK

**Author notes:** Co-corresponding author. Mailing address: 9000 Rockville Pike, Building 10, Rm 11C208, MSC 1888, Bethesda, MD 20892.

## Abstract

Human infection with *Cryptococcus* causes up to a quarter million AIDS-related deaths annually and is the most common cause of non-viral meningitis in the United States. As an opportunistic fungal pathogen, *C. neoformans* is distinguished by its ability to adapt to diverse host environments including plants, amoeba and mammals. In the present study, comparative transcriptomics of the fungus within human cerebrospinal fluid identified expression profiles representative of low-nutrient adaptive responses. Transcriptomics of fungal isolates from a cohort of HIV/AIDS patients identified a low nutrient-induced gene, an alternative carbon nutrient transporter *STL1* associated with poor early fungicidal activity, an important clinical prognostic marker. Mouse modeling and pathway analysis demonstrated a role for *STL1* in mammalian pathogenesis and revealed that *STL1* expression is regulated by a novel target-of-rapamycin (TOR)-related multi-gene regulatory mechanism involving the *CAC2* subunit of the chromatin assembly complex 1, CAF-1. In this pathway, the TOR-related RNA chaperone, *VAD1* was found to transcriptionally regulate a cryptococcal homolog of a cytosolic protein Ecm15, in turn, required for nuclear transport of the Cac2 protein. Derepression of *STL1* by the *CAC2*-containing CAF-1 complex was mediated by Cac2 and modulated binding and suppression of the *STL1* enhancer element. Derepression of *STL1* resulted in enhanced survival and growth of the fungus in the presence of low nutrient, alternative carbon sources, facilitating virulence in mice. The study underscores the utility of ex vivo expression profiling of fungal clinical isolates and provides fundamental genetic understanding of saprophyte adaption to the human host.

**Author summary:** The fungus *Cryptococcus* is a fungal pathogen that kills an estimated quarter of a million individuals yearly and is the most common cause of meningitis in the United States. The fungus is carried in about 10% of the adult population and, after re-activation, causes disease in a wide variety of individuals including HIV-infected as well as immunosuppression either from genetic defects or after immune suppressive treatments due to transplant conditioning, cancer therapy or treatment of autoimmune diseases. The fungus is widely carried in the soil and trees and can infect plants, single cell organisms and even dolphins. However, mechanisms for this widespread ability to infect a variety of hosts are poorly understood. The present study identified adaptation to low nutrients as a key property that allows the fungus to infect these diverse hosts and identified a nutrient transporter, *STL1* to be associated with a marker of poor clinical outcome in a cohort of HIV/AIDS patients. Understanding molecular mechanisms involved in environmental adaptation may help to design better methods of control and treatment of widely dispersed fungal pathogens such as *Cryptococcus*.

## Introduction

*Cryptococcus neoformans* (Cn) is a major fungal pathogen causing a highly lethal meningoencephalitis (CM) primarily in individuals with impaired host cell immunity such as those infected with HIV/AIDS, causing a quarter million deaths annually [1, 2]. Mortality exceeds 20-50% despite therapy [3], and recent failed attempts to improve outcomes [4] highlight our profound lack of understanding of the pathophysiology of human infections. While studies in host models such as mice or invertebrates have generated essential insights into fungal virulence, evolutionary differences in host response and pathogen environment suggest a need to model studies using human disease correlates [5].

Peculiar to *Cryptococcus* is the ability to live for extended periods both within the environment [6] and within the mammalian host in a dormant state [7]. It can also cause disease in a wide variety of plants [8], free living amoeba [9] and animals such as dolphins [10] and humans [11]. Similarities between mammalian host environments such as the macrophage phagolysosome and those more primitive such as amoeba have been implicated as exerting evolutionary pressure on facultative intracellular pathogens such as Cn [12] and gene expression studies suggest adaptation to nutrient deprivation is important within this environment [13]. Indeed, requirements for the gluconeogenesis enzyme Pck1 [14] and a high affinity glucose transporter [15] suggest roles for nutrient homeostasis in cryptococcal virulence. Phagocytosis also stimulates a starvation response in other pathogenic fungi such as *Candida albicans* that induce a shift to fatty acids as a carbon source by upregulating the glyoxylate cycle, requiring the enzyme isocitrate lyase [16]. However, the dispensability of isocitrate lyase by Cn during infection suggests that species-specific pathways for starvation tolerance are also important within the mammalian host [17]. In addition, induction of the nutrient-recycling autophagy pathway is particularly important for Cn but less important for other fungal pathogens such as *Aspergillus* and *Candida* [18]. For this important stress response, global nutrient regulators such as the target of rapamycin (TOR)[19] play an important link between cryptococcal starvation response, macrophage survival and pathogen fitness [20]. In addition, downstream of TOR the virulence-associated dead box protein, Vad1 has been shown to act as a global post-transcriptional regulator of TOR-dependent processes such as autophagy [20]. Similarly, the role of the cAMP nutrient sensing pathway has featured prominently in cryptococcal pathogenesis [21]. In this pathway, adenylyl cyclase is activated by a Ga subunit (Gpa1), resulting in the production of cAMP that binds to regulatory subunits Pkr1 of the PKA complex to release an active form of the catalytic subunit Pka1 which activates downstream proteins [22]. However, little is known regarding regulatory relationships that link these nutrient master regulators and gene targets responsible for mammalian virulence as well as possible commonalities between the various infective environments including humans.

Previous studies have not found robust relationships between mammalian infectious outcome and typical virulence factors in vitro, suggesting a need to identify new virulence attributes by novel approaches [23] [24]. Thus, to characterize virulence-associated starvation responses more relevant to human infections, transcriptional profiling of the fungus inoculated into human cerebrospinal fluid (CSF) was first compared to that from a starvation medium previously used to identify a clinical association between expression levels of a *CTR4* copper transporter and dissemination to the brain in a cohort of human solid organ transplant patients [25]. These studies demonstrated a high concordance between expression levels by the fungus within infected human cerebrospinal fluid and under nutrient deficiency and identified a large set of 855 upregulated genes. To identify genes within this subgroup whose expression might be most relevant to human infections, isolates from a cohort of HIV/AIDS-infected patients with CM were utilized in a discovery study of attributes related to 10 week mortality, along the lines of a previous hypothesis-driven analysis linking clinical outcome and known virulence attributes [24]. The second most highly differentially-expressed annotated gene (CNAG_01683/CP022331.1) showed highest homology to a <u>s</u>ugar transporter-like gene *STL1* from *Saccharomyces cerevisiae* implicated previously in glycerol metabolism. *STL1* expression levels showed a significant correlation with poor early fungicidal activity (EFA), a clinical prognostic marker of patient outcome [26]. Furthermore, mouse modeling demonstrated attenuated virulence of an *stl1*Δ strain, confirming its role in mammalian virulence. *STL1* was also identified as a target gene of TOR, regulated by a novel regulatory pathway involving mRNA decay and a multi-component Cac2-Cac1-Ecm15 nuclear shuttle regulatory complex. These studies thus provide a direct link between TOR signaling starvation response and CAF1-mediated regulation of microbial pathogenesis, relevant to human fungal infections.

## Results

### Transcriptional profiling of Cn suggest similarity in adaptive stress response between human CSF and starvation media at 37°C

Previous studies utilized nutrient-deficient media to simulate the environment of the host CSF and successfully correlated expression levels of a *CTR4* copper transporter with dissemination to brain in a cohort of solid organ transplant patients [25]. These conditions were utilized to compare transcriptional profiles of a pathogenic Cn strain (serotype A, H99), between nutrient rich and nutrient poor environment and compared the starvation adaptation profiles to that obtained after transfer to pooled human cerebrospinal fluid from patients with cryptococcal meningitis (S1 and S2 Table). A strong correlation was demonstrated between expression level changes in CSF vs. starvation media at 37°C (Fig 1A; Rsquare = 0.79, slope non-zero p < 0.0001). Transcriptional profiles after transfer to each respective condition demonstrated not only common identities of upregulated genes, but also the comparatively similar magnitudes of the transcriptional changes. Of 1,363 genes upregulated in starvation and 1,089 in CSF, 855 were upregulated in both conditions of which 332 were annotated sufficiently to allow GO term assignments (Fig 1B). Compared to the overall annotated transcriptome (Fig 1C, left panel), transition of Cn to either starvation conditions or CSF resulted in an expansion of gene expression associated with the gene ontology (GO) biological term “carbohydrate metabolic processes” including key enzymes involved in β-oxidation, the glyoxylate cycle and gluconeogenesis (S1 Fig). In contrast, genes in the category “translation” were under-represented in both media, consistent with the slow growth anticipated under these conditions. Using GO functional terms, transition of fungal cells to either starvation or CSF resulted in differential expression of genes involved in the transport function (S2 Fig). These data thus emphasize similarities in gene expression profiles of Cn between nutrient deprivation and CSF as well as a predominance of genes involved in carbohydrate homeostasis and transport function.

**Fig 1.**
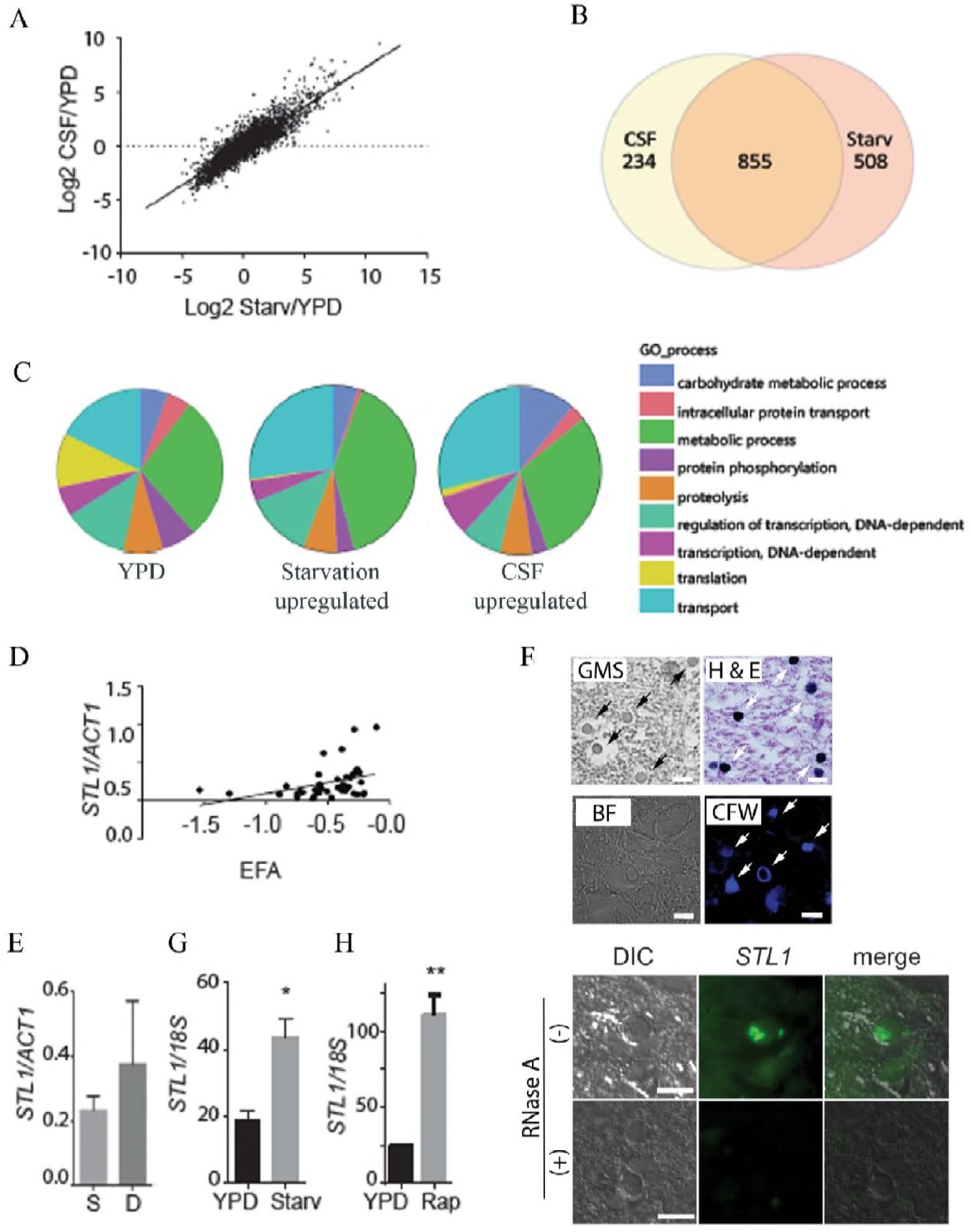
Expression profiling demonstrates a concordance of cryptococcal genes upregulated in fungal-infected human cerebrospinal fluid and starvation. (A) *C. neoformans* strain H99 was grown to mid log phase in YPD, then subjected to starvation in asparagine salts without glucose or sterile cerebrospinal fluid (CSF) at 37°C for 3 h. RNA was recovered and subjected to microarray analysis and values expressed as Log2 ratios of expression in CSF/YPD. N=2 for each condition. (B) Venn diagram of genes upregulated > 2-fold and adj. p < 0.05 after incubation in fungal infected CSF or under starvation conditions (Starv). (C) GO biological process term occurrence among genes upregulated under the indicated conditions. (D) Transcriptional ratio of *STL1/ACT1* determined by qRT-PCR of patient isolates compared to EFA (N = 46; regression analysis, N-2 d.f. p < 0.05. (E) Expression ratios of *STL1*/*ACT1* in survivors at 10 weeks (S, N=35) vs those who died (D, N=11). (F) Sections of a brain autopsy specimen were obtained from a 42 y.o. with HIV/AIDS who died of severe and diffuse *C. neoformans* infection and visualized by bright field microscopy (BF) or differential interference contrast microscopy (DIC) or stained with Gomori methenamine silver stain (GMS), haematoxylin and eosin stain (H&E stain), calcofluor white (CFW), or a C3-Fluoresein labelled oligonucleotide probes for *STL1* transcripts (*STL1*) by fluorescent in situ hybridization in the presence (+RNase A) or absence (-RNase A) RNase A treatment. Arrows point to stained yeast cells. Bar = 10 μm. (G, H) Expression levels of *STL1*/18S under mid-log growth in YPD or 3 h incubation under starvation conditions (Starv) or in the presence of 5 μM rapamycin (Rap). N = 3 independent experiments, Student’s t-test, *; p < 0.05; ** p < 0.01.

### Expression profiling of isolates from a cohort of AIDS-related CM identifies a putative <u>s</u>ugar <u>t</u>ransporter <u>li</u>ke gene, *STL1* as a candidate biomarker related to clinical outcome

Further studies were then conducted to determine whether starvation-induced genes may provide candidate biomarkers of clinical outcome that may help to understand human-related pathogenicity of the fungus. Transcriptional profiling under starvation conditions was conducted as described, which previously identified the copper transporter *CTR4* as a potential biomarker of brain dissemination in solid organ transplant recipients [27] using a set of serotype A Cn isolates from a cohort of HIV/AIDS patients described previously [28]. A demographic table of the cohort from which isolates were obtained is shown in Table 1. Median age was 36 and all were treated with amphotericin B-containing regimens. Despite therapy, 5 died by 2-weeks and an additional 6 died by 10-weeks, generating a 10-weeks mortality of 11/45 (24%), typical of CM in developing countries [3]. Pre-treatment co-variables were significantly different for mental status as measured by Glasgow coma score (p = 0.02), but parameters including CSF initial fungal burden, EFA, CSF opening pressure or WBCs or CD4 count did not differ significantly. Expression profiles were compared using ANOVA with adjustment, grouping patient isolates associated with mortality at 10 weeks vs. survivors. Ninety-one genes (28 annotated) showed significantly differing expression in patients who died vs. those who lived at least 10-weeks after therapy (S3 Table, adj p < 0.02). The first highly expressed gene was the nitroreductase family protein gene (CNAG_02692/CP022323.1). The second most highly differentially-expressed gene showed highest homology to a <u>s</u>ugar transporter-like gene *STL1* from *S. cerevisiae* previously implicated in glycerol transport [29, 30] and was selected for further study because of its putative role in alternative carbon metabolism and transport function exemplative of the aggregate transcriptional changes in human CSF. *STL1* expression levels by qRT-PCR from the clinical isolates showed a significant correlation with poor early fungicidal activity (EFA, Fig 1D; p = 0.018) [26] as well as a trend with clinical outcome (Fig 1E; p = 0.07); however, the latter was not statistically significant, possibly due to non-microbiological confounders such as immune responses or delays in medical care. EFA is the rate of fungal clearance in each of the HIV patient’s CSF during therapy and is a microbiological prognostic marker of clinical outcome [26, 31]. In addition, *STL1* expression was demonstrated in a brain autopsy specimen from an HIV/AIDS patient with CM using fluorescent in situ hybridization (66/66 yeast cells with +*STL1* signal vs. 9/78 in RNase-treated negative control slides, Fisher’s exact test, p < 0.0001; Fig 1F) [20]. Furthermore, qRT-PCR demonstrated that *STL1* expression was induced during starvation as well as in the presence of the TOR inhibitor rapamycin (Fig 1G, H). In summary, these data identify a starvation-induced potential biomarker of clinical outcome, *STL1*, associated with poor microbiological clearance (EFA) during HIV-associated human infections.

**Table 1.** Baseline clinical and laboratory characteristics

|  | All patients | Survived | Died |
| --- | --- | --- | --- |
| N | 45 | 34 | 11 |
| <sup>1</sup>Age (years) | 36 (21-62) | 37 (21-62) | 33 (23-51) |
| Sex (% male) | 53 | 53 | 55 |
| Abnormal mental status (%) | 9 | 3 | 27 |
| Fungal burden (log10 CFU/ml) | 5.9 | 5.9 | 5.8 |
| EFA | -0.51 | -0.55 | -0.37 |
| <sup>1</sup>CSF OP (cmH<sub>2</sub>O) | 26 (7-76) | 26 (10-76) | 15 (7-42) |

<sup>1</sup>median (range)
CSF, colony forming unit, CSF cerebrospinal fluid, EFA, early fungicidal activity, OP opening pressure

### Identification of a *VAD1*-dependent regulator of *STL1*, *ECM15*

Further studies were conducted to identify TOR-dependent regulatory pathways that could play a role in expression of starvation-associated genes such as *STL1*. TOR is a particularly important pathway as the TOR inhibitors sirolimus and everolimus, are in widespread clinical use as immunosuppressants [32]. Recently, the cryptococcal RNA chaperone Vad1 was shown to regulate gene expression in a TOR-dependent fashion by recruiting mRNA to a decapping complex protein Dcp2 for decapping, leading to transcript degradation [20]. Previously, we determined the mRNA binding profile of Vad1 in Cn by using RNA immunoprecipitation followed by microarray analysis (RIP-ChIP), and showed that *VAD1* binds multiple transcripts, including a gene *ECM15* [33]. *ECM15* in *S. cerevisiae* is a non-essential poorly understood protein, proposed to be involved in cell wall competency and was found on a yeast-hybrid screen to bind to a nuclear protein, Cac2 [34, 35]. In the current study, *ECM15* mRNA binding to Vad1 was confirmed by qRT-PCR after immunoprecipitation of a c-myc-Vad1 fusion protein compared to that of an equivalent precipitation using an untagged strain (Fig 2A). In addition, as shown in <u>Fig 2B</u>, a *vad1*Δ mutant showed accumulation of *ECM15* transcripts compared with wild-type from cells incubated under starvation conditions. Transcriptional profiling of an *ecm15*Δ strain showed reduced expression of the target gene, *STL1* described above (S4 Table), suggesting a regulatory circuit between TOR-dependent regulation and *STL1* through *VAD1* and *ECM15*. Phenotypic studies demonstrated a role for *ECM15* in expression of known virulence factors of *C. neoformans*. For example, the anti-phagocytic extracellular capsule was increased in the *ecm15*Δ mutant cells compared to wild-type under both nutrient replete conditions (Glu +) and nutrient deplete conditions (Glu -) which reverted in the complemented strain (Fig 2C). However, expression of the multifunctional laccase enzyme [24] was reduced in the *ecm15*Δ mutant (Fig 2D) as was mating (Fig 2E). These studies thus implicate a role for *ECM15* in the regulation of several virulence-associated phenotypes as well as mating of *C. neoformans*.

**Fig 2.**
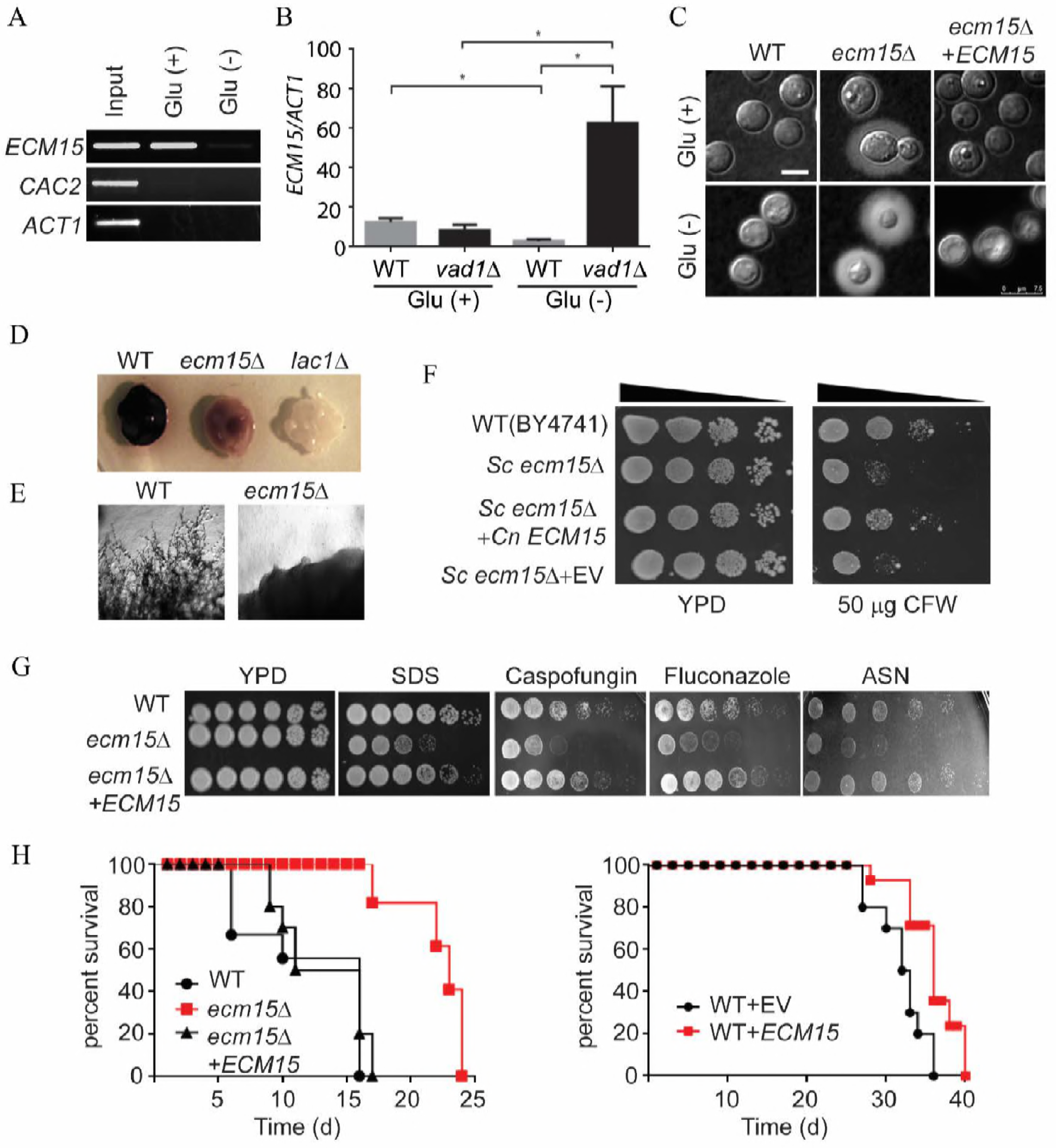
Deletion of cryptococcal *VAD1* increases transcript abundance of *ECM15*, a regulator of capsule, mating, laccase activity, and virulence. (A) Lysate from cells expressing a c-myc-tagged Vad1 fusion protein incubated under the indicated conditions were immunoprecipitated followed by RT-PCR/gel electrophoresis using primers for the indicated gene transcripts. (B) Steady state transcript levels of *ECM15* from indicated strains grown to mid-log in YPD (Glu +) and ASN salts without glucose (Glu -) media. N = 3 independent experiments, Student’s t-test, *; p < 0.05. (C) Indicated strains were incubated on YPD (Glu +) and ASN salts without glucose (Glu -) agar for 5 days at 30°C and examined by India ink microscopy. (D, E) The indicated cells were inoculated on ASN containing 100 mg/ml norepinephrine and observed after 2 days at 37°C (upper panel). WT and *ecm15*∆ mutant *MAT*α strains were co-incubated with a *MAT***a** mating partner (strain KN99) on nitrogen limiting mating media (V8 agar) for 2 weeks at 25°C. The edges of the mating mixtures were photographed (40x) (lower panel). (F) The indicated strains were diluted to an A_600_ of 1.0, and 1:5 serial dilutions (5 μl) were plated on YPD media or YPD containing the 50 μg Calcofluor White (CFW) and incubated at 30°C for 3 days. (G) The indicated strains were diluted to an A_600_ of 1.0, and 1:5 serial dilutions (5 μl) were plated on YPD media containing the indicated drugs and incubated at 30°C for 3 to 7 days. (H) Mice were inoculated by tail vein [10^6^ (left) or 10^3^ (right) of the indicated strains], and progress was followed until they were moribund. N = 10 mice per group, log ranks, (left panel p< 0.0001; right panel, p<0.01).

Further studies sought to investigate the functional relatedness of the cryptococcal Ecm15 protein to that of *S. cerevisiae*. The deduced amino acid sequence showed 66.8% identity to its respective homolog of Sc. Susceptibility of a Sc *ecm15*Δ mutant to the cell wall active agent Calcofluor white previously reported [36] was complemented by the cryptococcal *ECM15* gene (Fig 2F). These results suggest that Cn Ecm15 is a functional homolog of Sc Ecm15p. Extending these results to cell wall phenotypes associated with virulence, *ecm15*Δ mutants were found to exhibit increased susceptibility to the ionic detergent SDS as well as the anti-fungal cell wall agent caspofungin or the cell membrane disruptor drug fluconazole (Fig 2G). Interestingly the *ecm15*Δ mutant also displayed poor growth on nutrient poor agar consisting of asparagine salts (ASN) without glucose consistent with its TOR dependence (Fig 2G). In addition, the Cn *ecm15*Δ strain exhibited attenuated virulence in a mouse model (Fig 2H). Interestingly attenuated virulence was also exhibited by an *ECM15* overexpressing strain compared to an identical strain containing an equivalent empty vector at the same copy number (right panel). ECM15 was overexpressed using an *ACT1* constitutive promoter and compared with strains expressing empty vector alone in equivalent copy number as previously described [37]. Attenuated virulence with deletion or overexpression is a hallmark of the cryptococcal *VAD1* gene that shows a similar phenotype [20] and may be due to their close regulatory association. Taken together, these data suggest that deletion of cryptococcal *VAD1* increases transcript abundance of *ECM15*, a regulator of capsule, mating, laccase, cell wall stability, starvation tolerance and mammalian virulence.

### *ECM15* dependent regulation of a chromatin assembly factor, *CAC2* involved in extracellular capsule expression and mammalian virulence

Previous whole proteome interaction studies suggested that Sc Ecm15p may interact with Sc Cac2 [34]. Cac2 is a key constituent of the CAF-1 chromatin assembly factor which assembles histones H3 and H4 and mediates chromatin suppression of genes at sub-telomeric locations and tolerance to ultraviolet irradiation (UV), the latter demonstrated in both Sc and Cn [38, 39]. However, its precise role as a potential regulator remains unexplored in eukaryotes. Construction of a Cn *cac2*Δ mutant confirmed UV sensitivity (Fig 3A) as reported previously in Sc and allowed a transcriptional comparison with the *ecm15*Δ mutant constructed in the same Cn genetic background. Deletion of *ECM15* resulted in at least a 2-fold increased transcription (Fig 3B; adj. p < 0.05) of 106 genes and *CAC2* in 8 genes, 4 shared with *ECM15*. Interestingly, the gene showing highest suppression by both *ECM15* and *CAC2* was *STL1* (S4 Table). Further study identified several virulence-related phenotypes shared between *CAC2* and *ECM15*. As shown in Fig 3C and S9 Fig, capsule sizes under nutrient deplete conditions were accentuated in the *cac2*Δ mutant (a phenocopy of the *ecm15*Δ mutant), and was suppressed to that of wild-type in the *CAC2* complemented strain. Epistatic studies demonstrated that capsule sizes under both nutrient replete and nutrient deplete conditions were restored to WT in *ecm15*Δ mutant cells overexpressing the *CAC2* gene, although overexpression of *ECM15* had no effect on the capsular phenotypes of the *cac2*Δ mutant cells. These results suggest that *CAC2* is a suppressor of capsule formation downstream of *ECM15*. In addition, *CAC2* transcription was suppressed in the *ecm15*Δ mutant strain under starvation conditions, suggesting that *ECM15* partially activates *CAC2* signaling through a transcriptional process. In addition, demonstration of *CAC2* transcript accumulation in the *vad1*Δ mutant, similar to *ECM15* suggests the strong effects *VAD1* as a global regulator (Fig 3D and 3E) [14]. Furthermore, *CAC2* overexpression resulted in suppression of virulence (Fig 3F), similar to that of the *ECM15* overexpressor strain, suggesting a role of both in the suppression of virulence. However, deletion of *CAC2* downstream of *ECM15* did not increase mammalian virulence suggesting loss of *ECM15*-regulated processes as the pathway was traced downward. Taken together, these data suggest an epistatic relationship between *ECM15* and *CAC2* in the repression of cryptococcal capsule and mammalian virulence.

**Fig 3.**
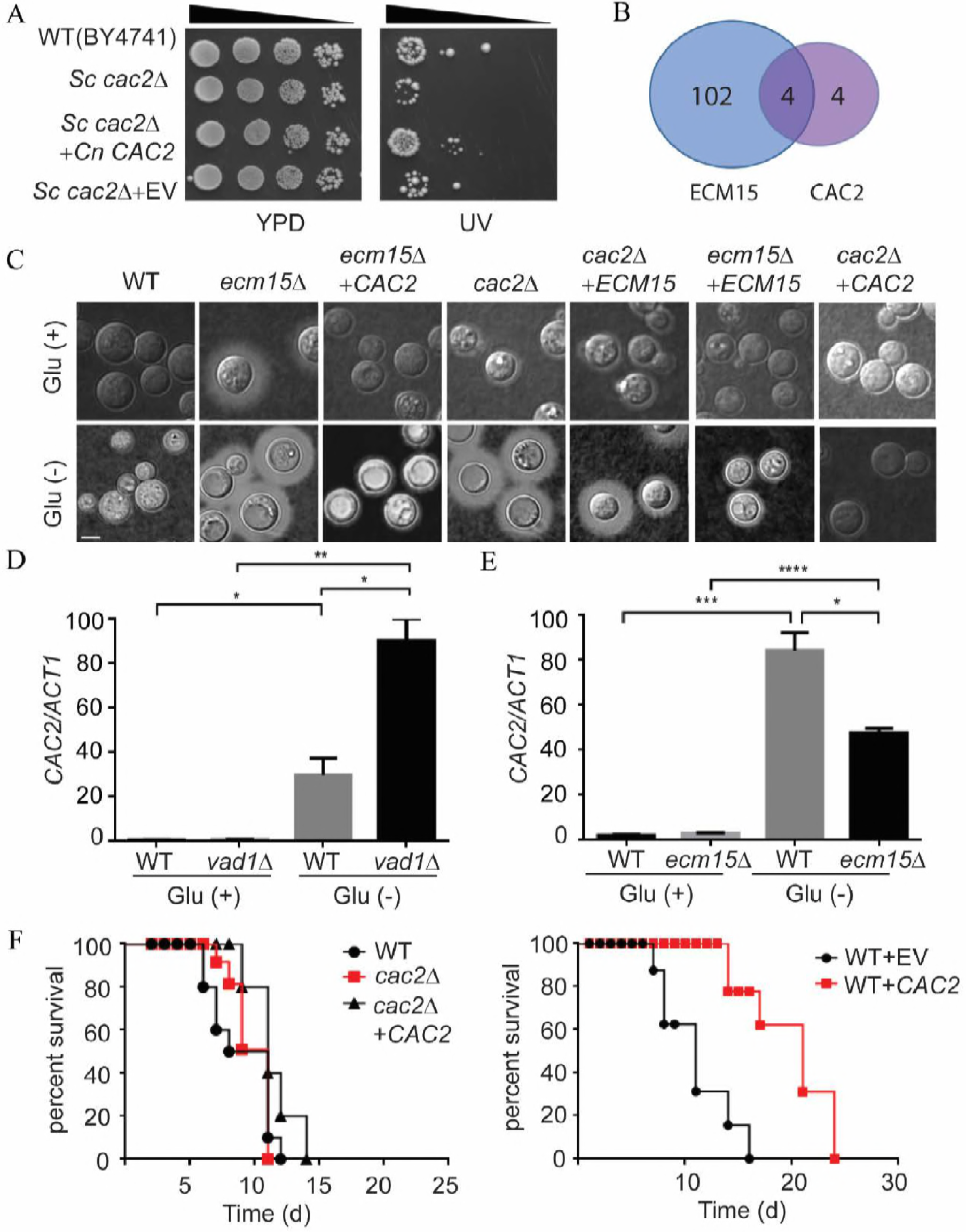
*ECM15* represses capsule formation via *CAC2*. (A) The indicated strains were diluted to an A_600_ of 1.0, and 1:5 serial dilutions (5 μl) were plated on YPD media or UV irradiation for 1 min and incubated at 30°C for 3 days. (B) Venn diagram of the number of genes showing >2x increased transcription relative to WT and adj p < 0.05 after deletion of *ECM15* or *CAC2.* (C) Indicated cells were incubated on YPD (Glu +) and ASN without glucose (Glu -) agar for 5 days at 30°C and examined by India ink microscopy. Bar = 5 μm. (D, E) qRT-PCR of *CAC2* of indicated strains grown to mid-log in YPD (Glu +) and ASN without glucose (Glu -) media (N = 3 independent experiments, Student’s t-test; *; p < 0.05, **; p<0.005, ***; p<0.0005, ****; p<0.0001). (F) Mice were inoculated by tail vein (10^6^ of the indicated strains), and progress was followed until moribund. N=10 mice per group, log ranks, left panel p=0.7; right panel: p < 0.0001.

### Nuclear localization of *CAC2* is nutrient-dependent and requires *ECM15*

*Saccharomyces* interactome studies [34], suggested that *ECM15* may regulate *CAC2* by a post-translational interactive mechanism. Expression of fluorescent-tagged Cac2 and Ecm15 fusion proteins demonstrated co-localization to the nucleus (Fig 4A) under nutrient rich conditions with a cytoplasm localization under starvation or in the present of rapamycin conditions, consistent with a possible role as nuclear suppressors under the former conditions and derepression under starvation conditions. Examination of proteins sequences using cNLS-Mapper [40] identified putative nuclear localization sequences in Ecm15 in the region D63-A91and Cac2 in the region E778-V802 (S3 Fig). Further studies demonstrated that deletion of *ECM15* resulted in mislocalization of the Cac2 fusion protein under glucose replete conditions or rapamycin treated conditions (Fig 4B). Interestingly, Ecm15 also mis-localized to the cytoplasm in *cac2*∆ mutant suggesting that an interaction between the two is required for effective nuclear co-localization. These results suggest that Ecm15 and Cac2 play an interacting role in nuclear targeting, leading to downstream repression of virulence associated phenotypes and genes such as *STL1* described in Fig 3.

**Fig 4.**
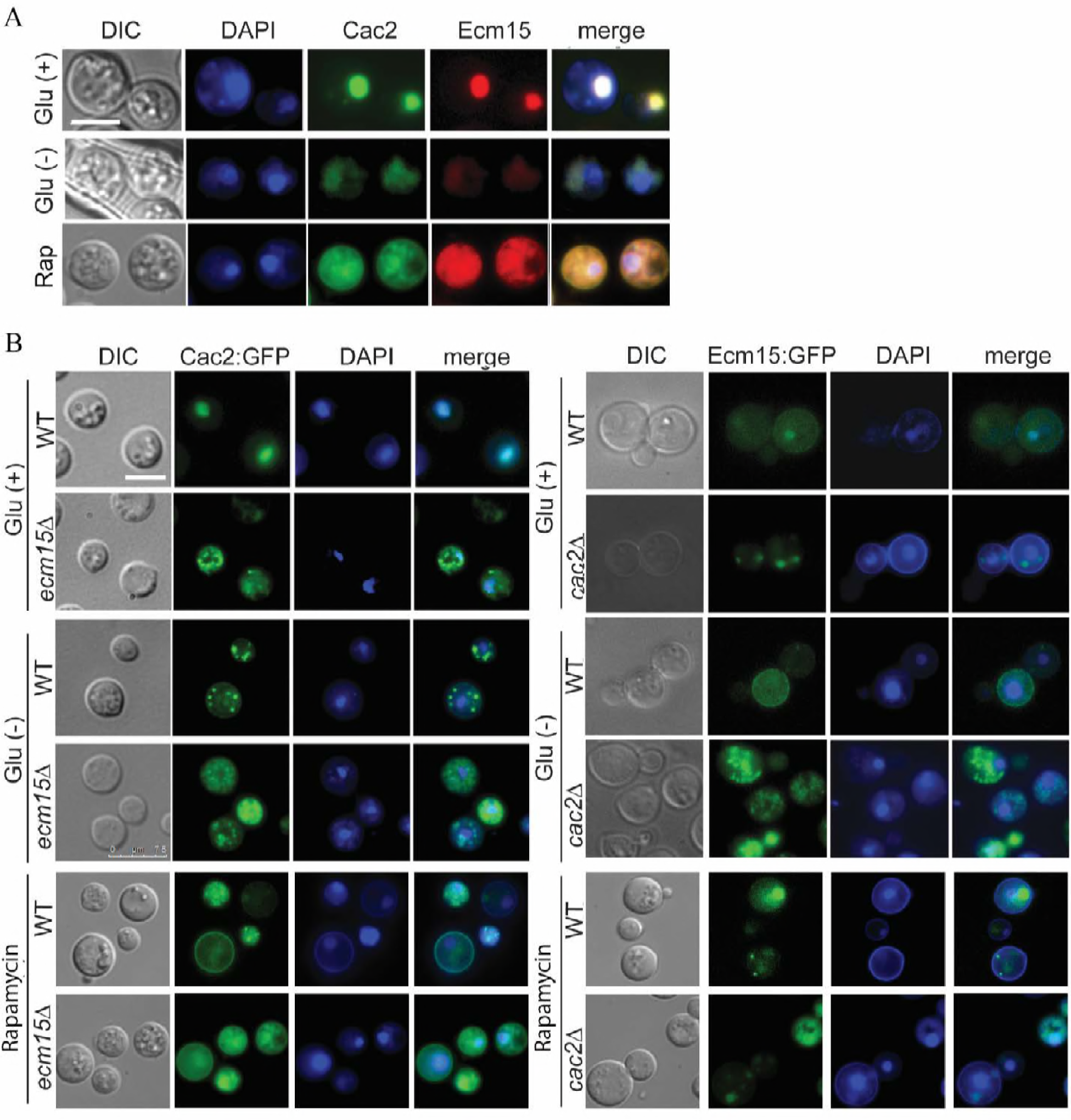
Cac2 localized by Ecm15. (A) *C. neoformans* cells expressing Cac2-GFP and Ecm15-mCherry were incubated on YPD (Glu +), ASN without glucose (Glu -) media and YPD containing rapamycin, and observed by fluorescence microscopy (DAPI, Cac2, Ecm15) or differential interference microscopy (DIC). Bar = 5 μm. (B) Cells expressing Cac2-GFP in wild-type and *ecm15*∆ were incubated on YPD (Glu +), ASN without glucose (Glu -) and YPD containing rapamycin, and observed for fluorescence or differential interference microscopy (DIC) (upper panel). Cells expressing Ecm15-GFP in wild-type and *cac2*∆ were incubated on YPD (Glu +), ASN without glucose (Glu -) media and YPD containing rapamycin, and observed for fluorescence or DIC (lower panel). Bar = 5 μm.

### *STL1* is negatively regulated by *CAC2* and a *stl1*∆ mutant has attenuated virulence in a mouse model

To confirm a role for *CAC2* in *STL1* regulation, qRT-PCR was utilized which demonstrated increased *STL1* expression in WT cells under starvation conditions or in *cac2*Δ cells under both nutrient and starvation conditions consistent with a role as an *STL1* repressor (Fig 5A). Increased wild-type expression of *STL1* under starvation conditions was likely due to less nuclear localization of *CAC2* demonstrated in Fig 4A. Further studies demonstrated a role for *STL1* in expression of the virulence factors capsule (Fig 5B), laccase (Fig 5C), and mating (S8 Fig). Because of its putative role in alternative carbon homeostasis and virulence in the clinical isolates, we tested to see if *STL1* overexpression could facilitate growth on alternative carbon sources. Interestingly, *STL1* overexpression facilitated increased growth in nutrient limitation media containing 0.03% of either pyruvate, lactate or acetate--intermediates in gluconeogenesis and previously identified substrates within mammalian brains during CM infections as well as glycerol, which has been previously described for the *STL1* homolog from *Candida* (Fig 5D) [29, 41]. In addition, *STL1* overexpression facilitated increased growth in nutrient limitation media containing either 0.03% of ribose, or citrate but not equivalent concentrations of glucose vs. an identical strain transformed with empty vector alone in identical copy number (Fig 5E). These data extend the spectrum of alternative carbon substrates related to Stl1. However, *STL1* did not facilitate growth on other sugars including galactose, glucosamine, and rhamnose, suggesting carbon substrate specificity (S4 Fig). In addition, the *stl1*Δ strain exhibited moderately reduced virulence in an intravenous mouse brain dissemination model compared to the wild-type strain (Fig 5F, left panel). However, inoculation of *Cn* strains overexpressing *STL1* resulted in no additional increase in virulence in the highly virulent strain H99, compared with identical strains transformed with empty vector alone (Fig 5F, right panel). Taken together, these data suggest that deletion of cryptococcal *CAC2* increases transcript abundance of *STL1*, a regulator of capsule, sugar transport, laccase activity, and virulence.

**Fig 5.**
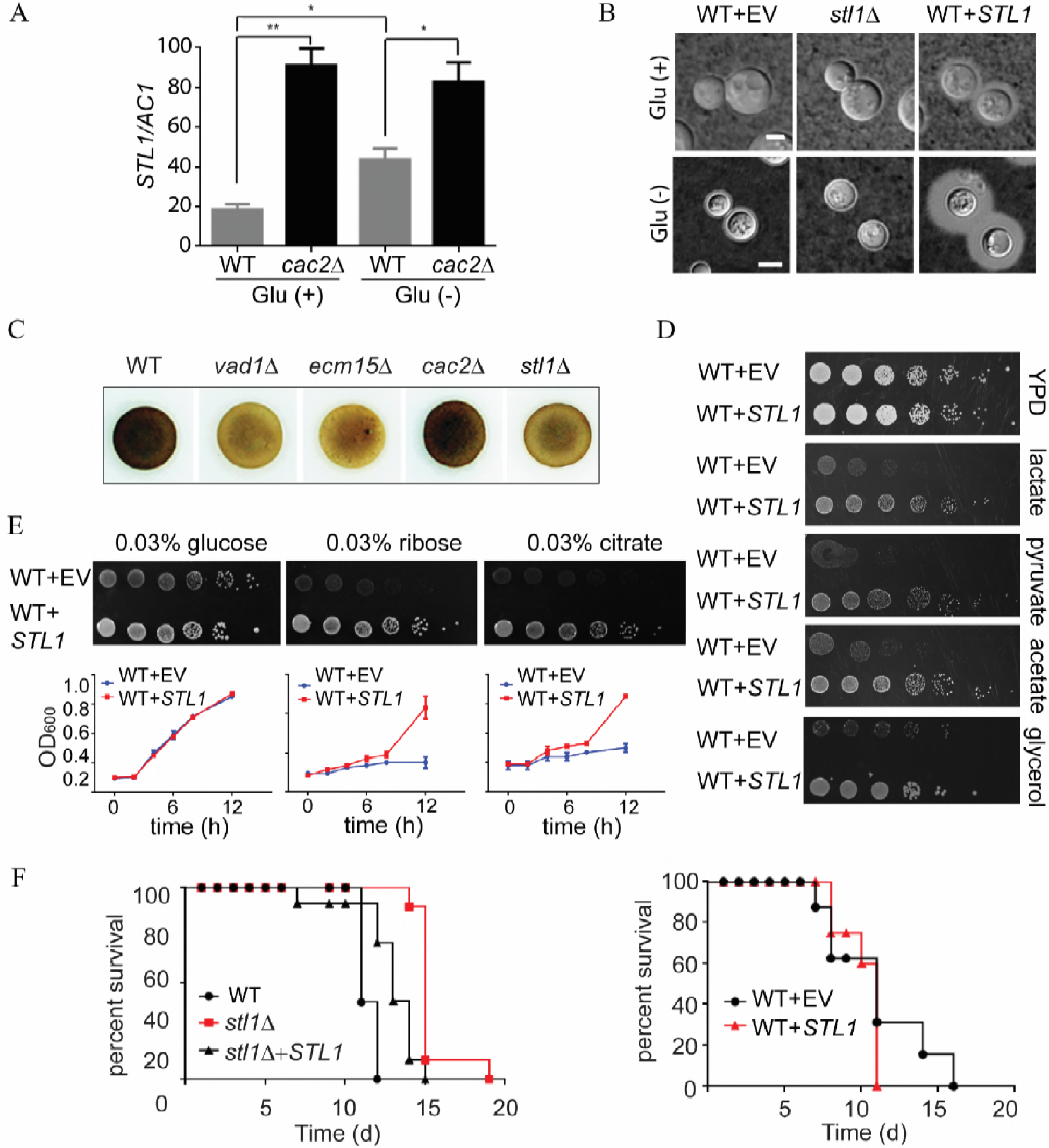
*STL1* is negatively regulated by *CAC2* and required for growth in alternative substrates, capsule formation, laccase activity and mammalian virulence. (A) qRT-PCR of *STL1* of indicated strains grown to mid-log in YPD (Glu +) and ASN without glucose (Glu -) media. N = 3 independent experiments, Student’s t-test, *; p < 0.05, **; p<0.01). (B) Indicated strains were incubated on YPD (Glu +), ASN without glucose (Glu -) agar for 3 days at 30°C and examined by India ink microscopy. Bar = 5 μm. (C) Indicated cells were inoculated on ASN containing 100 mg/ml norepinephrine to assay production of melanin pigment by laccase. (D) The indicated strains were diluted to an A_600_ of 1.0, and 1:5 serial dilutions (5 μl) were plated on YPD media or ASN containing the indicated substrates (0.03%) and incubated at 30°C for 5 days. (E) The indicated strains were diluted to an A_600_ of 1.0, and 1:5 serial dilutions (5 μl) were plated on ASN media containing the indicated substrates (0.03%) and incubated at 30°C for 7 days (upper panel). Indicated strains were cultured with ASN broth containing the indicated substrates (0.03%), and optical density (OD_600_) were calculated in indicated time points (lower panel). (F) Mice were inoculated by tail vein (10^6^ of the indicated cells), and progress was followed until moribund. N = 10 mice per group, log ranks, (left panel: p < 0.0001; right panel: p=0.5).

### Cac2 exhibits TOR-dependent *STL1* promoter occupancy

Early studies suggested that orthologs of CAF-1 members such as Cac2 are associated with suppression of genes in the sub-telomeric region [38], which include cryptococcal genes such as *FRE7* (CNAG_00876/ CP022325.1), CNAG_05333/XM_012198174.1 and the HpcH/Hpa1 (CNAG_06874/XM_012194223.1) family protein gene (S4 Table); however, the *STL1* gene does not reside within this region (location; chromosome 11, CP003830.1: 603,867-606,604). Thus, to identify possible direct *STL1* Cac2-specific DNA binding region(s), chromatin immunoprecipitation (ChIP) was performed which demonstrated promoter occupancy within a 100-bp fragment centered at −700 bp from the transcriptional start site of *STL1* as well as that of *FRE7* in nutrient condition (Fig 6A), otherwise, *STL1* Cac2-specific DNA binding efficiency was reduced in both starvation and rapamycin condition (S5 Fig). To determine the functional significance of the binding, we tested expression levels of a Stl1-green fluorescent protein (GFP) fusion having serial deletions of the *STL1* promoter (−1000, −500, −50 from the transcript start site) on basal transcriptional activity in *C. neoformans*. The plasmid containing an *STL1* ORF with various lengths of the *STL1* 5′-promoter GFP fusion gene were transformed into wild-type or *cac2*Δ *C. neoformans* strains. Empty vector was used as a control. These results indicated the presence of basal transcription originating from the −1,000 to −500 region of *STL1* under both nutrient replete (Glu +) and nutrient poor (Glu -), corresponding to the region of Cac2 binding by ChIP (Fig 6B and 6C). However, in the nutrient-rich condition, *CAC2*-dependent activity in the WT strain was only partially suppressed using the episomal constructs (S6A and S6B Fig) and suggested a requirement for additional chromatin-dependent promoter-binding factor(s) for effective *CAC2*-dependent chromatin silencing [42]. Thus, linear PCR fragments containing the indicated promoter, *STL1* coding region and selection marker were transformed into WT and *cac2*Δ strains as integrated fragments. As shown in Fig 6B and 6C, the integrated construct exhibited more strongly repressed *STL1* expression under nutrient rich conditions (Glu +), which was derepressed in either nutrient poor (Glu -) or in the *cac2*Δ mutant strain (S6 Fig). Transcriptional studies by qRT-PCR confirmed the *CAC2*-dependent suppression (S7 Fig). Interesting was the heterogeneity in the expression of the *cac2*Δ strain which may suggest additional post-translational regulatory features.

**Fig 6.**
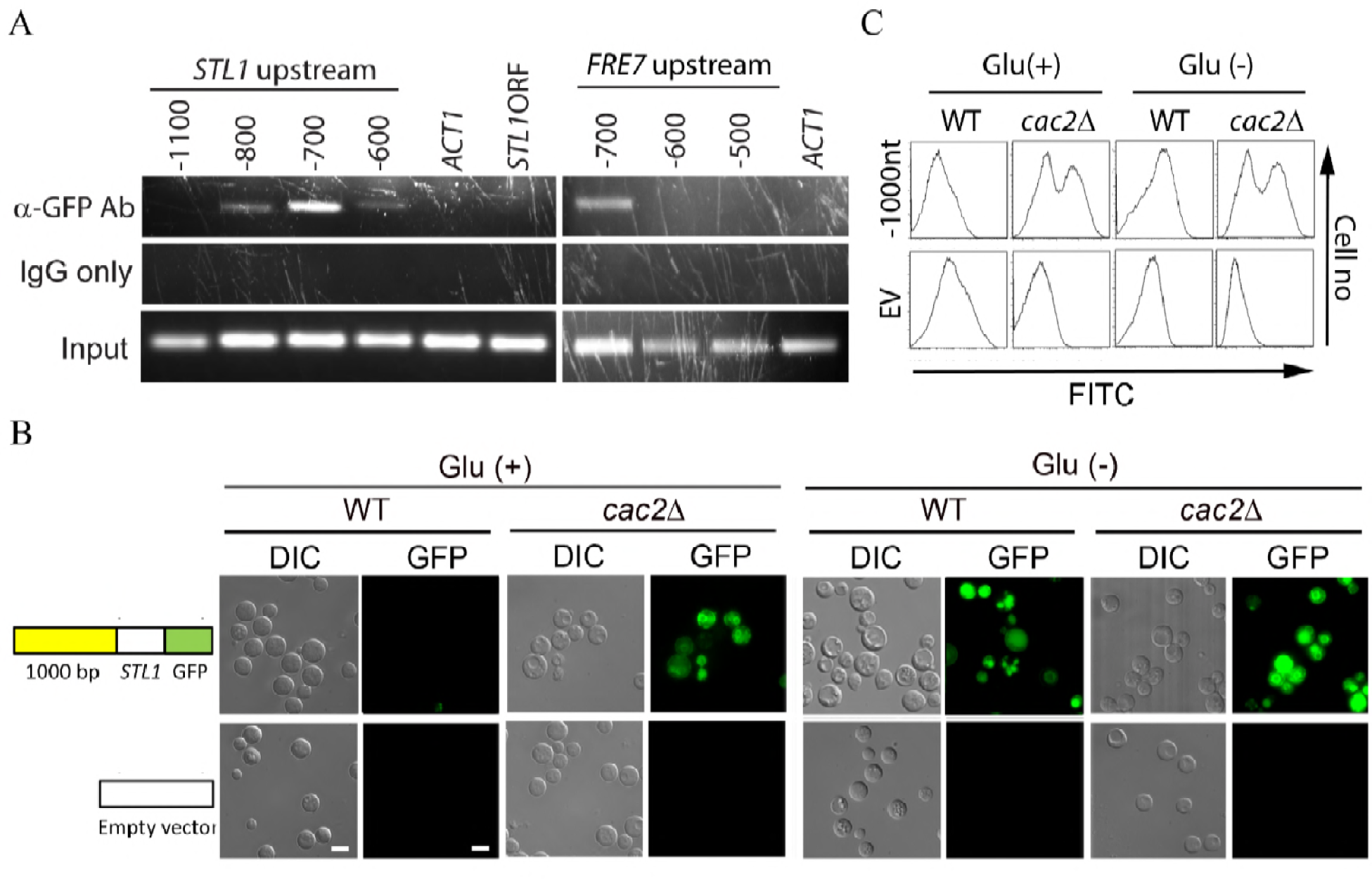
Chromatin immunoprecipitation (ChIP) survey of Cac2-promoter complexes. (A) Chromatin immunoprecipitation (ChIP) of a Cac2-promoter complex. Nuclear extract from induced cryptococcal cells was immunoprecipitated as described in Methods and assayed by PCR for the presence of the indicated regions of *STL1* or *FRE7* promoter sequences or control promoter sequences of *ACT1*. (B, C) Effects of serial deletion of the *STL1* 5’-promoter (−1000 from the transcript start site) on basal transcriptional activity in *C. neoformans*. Linear constructs containing *STL1* ORF with indicated regions of the *STL1* 5′-promoter GFP fusion gene were transformed into WT or *cac2*Δ strains of *C. neoformans* and grown to mid-log in YPD (Glu +) and ASN salts without glucose (Glu -) media and the population subjected to microscopy (DIC, GFP fluorescence) or flow cytometry (FITC). Empty vector used as a control. Bar=5μm.

In summary, these data identify a TOR-dependent epistatic regulatory pathway involving *VAD1*, *ECM15* and *CAC2* controlling growth on alternative carbon sources, expression of the virulence factors capsule and laccase and mammalian virulence (Fig 7).

**Fig 7.**
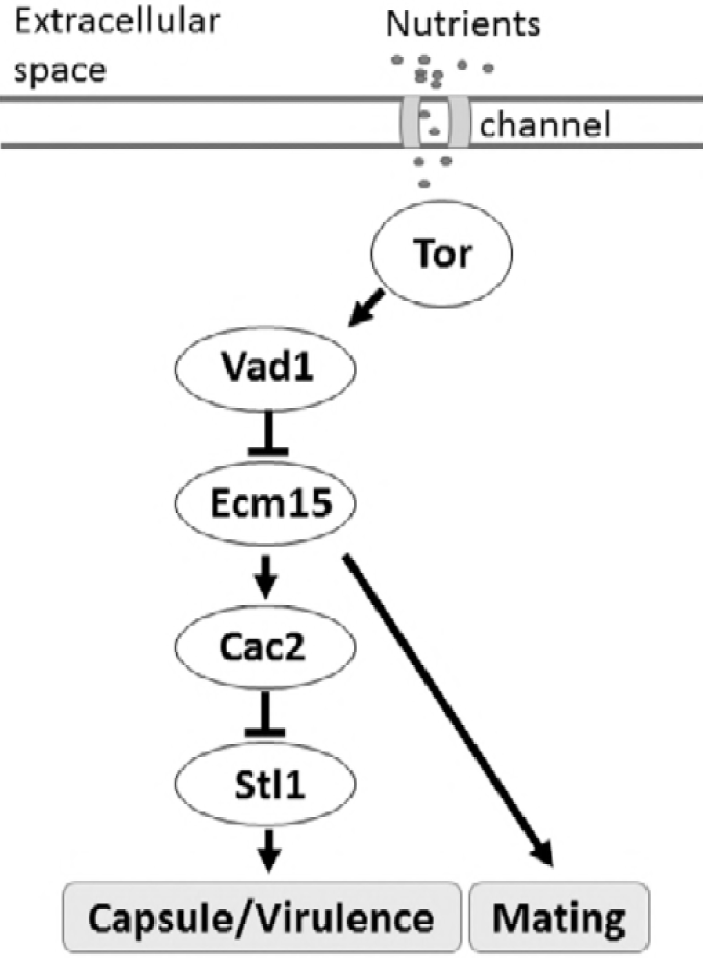
Scheme of TOR regulation of *STL1* related to capsule formation, and mammalian virulence.

## Discussion

Pathogen survival and virulence entails selection for traits necessary within the infective niche. For opportunist pathogens such as *C. neoformans*, evolutionary pressure exerted during environmental residence by resistance in free-living amoeba [9] or in the presence of diphenolic toxins within plants [8], have likely shaped the expression patterns necessary for effective mammalian infection. In the example of *C. neoformans*, the organism is also thought to reside in the host in many cases as a latent organism, which undergoes reactivation after immunosuppression by diseases such as HIV [7]. Thus, expression patterns of environmental *C. neoformans* strains may also undergo optimization through microevolution after exposure to the mammalian infective niche which has been demonstrated in mice [37] and may influence clinical outcome in humans [25]. These combined pressures have resulted in an organism highly adapted for survival and growth under nutrient limitation conditions [43]. The molecular patterns that shape adaptation and optimization within these different environments was the subject of the present study.

In these studies, expression patterns of *C. neoformans* within human cerebrospinal fluid from actively infected patients was highly correlated with that induced under starvation conditions at 37°C both in identity and levels of transcription. In addition, of the 222 genes previously reported to be upregulated after incubation in the macrophage cell line J774.1 [13], 147 were also found to be upregulated in CSF in the present studies and of the 323 genes upregulated in free living amoeba reported in the same paper, 164 were also upregulated in CSF and similar numbers in starvation in the present studies, suggesting metabolic response commonality between these infective niches. Brain infections with *C. neoformans* are typically associated with low CSF glucose levels [3] and the pooled patient CSF reflected this, ranging from 10-25 mg/dL (0.55-1.4 mmol/l) in the present studies. Interestingly, several studies suggest that the macrophage phagolysosome is also a nutrient-deprived environment and such stress results in pathogen adaptation for survival [4, 37]. These commonalities reflective of starvation response thus implicate starvation response pathways in cryptococcal virulence in humans. However, these expression patterns differed from that reported from RNA isolated directly from patient CSF of two CM patients that more closely resembled that from nutrient replete media and could have reflected an unusually vigorous growth condition or higher levels of CSF glucose concentrations typical of some HIV/AIDS patients [44]. However, such results caution that expression within a number of different environments may be required to understand heterogeneity within the human infective niche. In the present studies, CSF incubation resulted in reduced expression of genes related to protein translation typical of growth limitations from reduced nutrients [45]. Conversely, exposure to infected human CSF resulted in increased gene expression of processes related to sugar transport and alternative substrate utilization exemplified by genes involved in β-oxidation, the glyoxylate cycle and gluconeogenesis. This metabolic reprogramming towards more efficient utilization of alternative substrates and low glucose is shared by other organisms such as *Candida albicans* that also modifies its metabolism to assimilate these alternative substrates [46]. However, increased expression alone does not necessarily identify genes important to virulence, exemplified by the virulence factor laccase that is down regulated at elevated temperatures [47]. Disappointingly, genes demonstrating increased expression within pathogenic niches have shown variable roles in mammalian virulence. For example, the highly upregulated pyruvate kinase *PYK1* and hexose kinase I and II (*HXK1*/*HXK2*) involved in alternative substrate demonstrate attenuated virulence in a mouse model [15], whereas the highly upregulated *PTP1* sugar transporter demonstrated no such role [13]. Other upregulated genes such as *PCK1* have had variable roles in virulence, with mutants attenuated in mice [14], but not in rabbits [15]. Such discordant results demonstrate the limitations of exclusive reliance on model host studies of virulence.

Thus, the present studies utilized multiple patient isolates from a cohort of previously described HIV/AIDS patients presenting with cryptococcal meningoencephalitis and treated uniformly with amphotericin-based regimens to study gene expression under the same nutrient limitation conditions described above. An important benefit of human-associated studies is that they may reflect better the specific environment of the human host vs. those derived from mouse studies alone, which may have limitations [48]. For example, mice strains such as C57BL/6J may elicit a significant neutrophilic and even eosinophilic response in lungs after infection with clinical fungal isolates, which is quite different from the histiocytic response of humans including giant cell formation, depending on the relative cellular immunity of the infected patient [49]. Drawbacks are that it is difficult to control for heterogeneity among unrelated fungal organisms and their hosts. The present studies identified a potential role for an *STL1* sugar transporter required for survival and growth under low nutrient conditions and was also prioritized for further study because it was representative of an alternative substrate and transporter gene population showing an expansion in expression after transition to infected human CSF. *STL1* is named as a sugar transporter but has been better characterized as having a role in glycerol acquisition [29, 50] which was also found in the present studies in addition to roles in lactate, pyruvate and acetate metabolism. Interestingly, glycerol is elevated during cell membrane degradation and brain injury [51] and acetate has been demonstrated in large amounts by magnetic imaging spectroscopy during human cryptococcal infections [52, 53] and thus *STL1* may play a role in acquisition of these metabolites during infection. Expression levels of *STL1* showed a trend towards and association with 10-weeks mortality in HIV-related CM and were also associated with an important prognostic marker of microbiological clearance during human infections, EFA. EFA is a research measure of CSF microbiological clearance in patients which reflects both microbiological and drug responses as well as host aspects of the infection that affects clearance and demonstrates reduced clearance in groups who die [54]. In addition to its role as a prognostic marker EFA has been secondarily proposed as a treatment surrogate, related to antifungal activities that distinguished amphotericin b vs. standard fluconazole therapy [26] but has not proven to be robust surrogate for the second role in multiple randomized-controlled studies of amphotericin-based regimens, according to Institute of Medicine guidelines [31]. Indeed, we found that EFA in the present cohort of patients treated with amphotericin-related regimens was broadly similar. Notable was the reduced expression of the virulence-factor laccase in the *stl1*∆ mutant which has previously been shown to be correlated with EFA and patient survival and could have contributed to microbiological retention in the CSF [24]. However, we were not able to demonstrate a statistically significant correlation of *STL1* expression with patient survival with the numbers of available patient isolates which may be due to the large variability in expression levels between wild-type strains. Such variability was previously demonstrated by other cryptococcal biomarkers such as *CTR4* which demonstrated over 100-fold difference among clinical isolates from a cohort of solid organ transport patients [25]. However, it would be premature to speculate as to the role of *STL1* as a clinical biomarker and is likely co-related to other microbiological markers. There also exists a number of potential non-microbiological factors that affect mortality. For example, initial patient mental status was more frequently altered in those who died and is an important risk factor for death [3], as was found in our cohort. Other previously reported clinical risk factors such as age and opening pressure on lumbar puncture did not differ between groups. Another important contributor to death in primary infections with *C. neoformans* is the relative immune response of the host although previous studies have shown only minor relationships of markers such as CD4 count, peripheral white blood count or CSF white count [3]. The present cohort did not include patients with cryptococcal immune reconstitution syndrome (cIRIS) where immune responses may be more predominant [55] or other inflammatory syndromes associated with non-HIV infected individuals [56]. All of these considerations led us to attempt to provide additional validation of the human cohort fungal expression studies by utilizing a mouse model which showed a modest difference in virulence between wild-type and *stl1*Δ strains, further suggesting a role for *STL1* in mammalian infections. The intravenous model was tailored to the clinical cohort that all had brain infections and focuses on pathogen-related outcomes of murine brain infection, rather than a pulmonary (intratracheal or posterior pharyngeal inoculation) model where survival is more related to altered lung pathology [57]. However, additional cohorts reflecting differing clinical scenarios may be required to fully understand the complexity of the cryptococcal-human host interaction.

The present studies also identified an important molecular pathway in starvation response and its relation to human infections by examining *STL1* as a target gene of the TOR stress pathway. TOR is an important mediator of the starvation/stress response, is important for cryptococcal survival [58] and inhibitors of TOR such as sirolimus and everolimus are in widespread use in patient populations at risk for CM including transplant recipients [59]. More recent work has shown that many TOR-dependent starvation processes are regulated via mRNA stability in yeast [60] and an important TOR-dependent regulator of mRNA stability, *VAD1* is also a major virulence determinant in *C. neoformans* [20]. The present studies identified a role for TOR/*VAD1*-dependent regulation of a novel *CAC2/ECM15* regulatory pair of nuclear factors that demonstrated starvation-dependent regulation of *STL1*. In *C. neoformans*, a deletant mutant of *CAC2* was previously found to exhibit normal growth characteristics and a slight susceptibility to UV irradiation [61]. In *S. cerevisiae*, deletion of CAF-1 subunits such as *CAC2* results in increased UV sensitivity and silencing defects of sub-telomeric genes [38], suggesting a role in stress response, but the present studies are the first describing the role of Cac2 as a dynamic regulator, per se. Interestingly, only *CAC2* overexpression resulted in reduced virulence in mice, but this would be expected as it is an *STL1* repressor. The virulence phenotype was more evident in the *CAC2* overexpressor than the *STL1* knockout mutant and suggests a combined effect on virulence of multiple *CAC2*-regulated genes that may not have reached significance in the patient cohort. Another *ECM15*/*CAC2* dependent gene, CNAG_00876/CP022325.1, has been previously identified as a ferric-chelate reductase (*FRE7*), regulating the important iron acquisition pathway in *C. neoformans* [62]. The *FRE7* gene was also upregulated in the patient cohort, though did not approach significance and was not further characterized. This regulatory pathway has been simplified into a linear chain for study purposes but is likely much more complex due to the presence of other known virulence pathways related to alternative carbon acquisition/glycerol acquisition and metabolism including *HOG1*, calcineurin, *PI3K*, *PLC1* and *PKA1* [20, 63-69]. Repression by *CAC2* may be chromatin dependent as promotor studies demonstrated greater *CAC2*-dependent suppression in glucose when the constructs were integrated. Indeed, previous studies suggested that the Cac2-Caf1 complex binds preferentially to modified histones that have not been reported in plasmid DNA [70] and may thus be required as part of the chromatin repressor complex [71]. The related factor Cac1 has recently been shown by single-particle electron microscopy to act as a histone binding platform, linking Cac2 within the Caf1-histone assembly complex implicated in maintenance of molecular architecture [72]; however, the present studies extend the role of Cac2 within the Caf1 complex from merely a house-keeping function to that of a dynamic regulator of TOR-mediated nutrient stress response. Future studies may further help to provide more detailed structural mechanistic insight for this complex regulation in eukaryotes and highlights the utility of *C. neoformans* as a model eukaryote. Nevertheless, the present data demonstrates that the *VAD1/ECM15/CAC2* regulatory pathway is integral to the connection of the starvation/TOR virulence response through the sugar transporter *STL1* (Fig 7).

## Methods

### Ethics Statement

Written informed consent was obtained and the study was approved by the Research Ethics Committee of the University of Cape Town, the Medicines Control Council of South Africa, and the London-Surrey Borders Research Ethics Committee on behalf of St. George’s University of London and by an institutional review board (IRB)-approved protocol from the National Institute of Allergy and Infectious Diseases. All experimental procedures were conducted under a protocol approved by the Institutional Animal Care and Use Committee of the Intramural Research Program of the NIAID, NIH (Protocol No: LCIM12E). All experimental studies were approved by the relevant NIAID Animal Care and Use committee, as per the “Laboratory Animals: For The Care And Use Of laboratory animals,” National Research Council of the National Academies, Washington, DC.

### Study Subjects

Subjects providing isolates for expression analysis were control participants in a randomized trial of adjunctive IFN-γ in HIV-infected patients, and a randomized trial examining alternative amphotericin B combinations, both described previously [73]. We attempted to standardize the antifungal drug factor/ impact on EFA by ONLY selecting for inclusion patients from a single clinical trial site treated using similar study protocols with AmB-based induction regimens (ie no fluconazole and no adjunctive IFN gamma both of which we know lead to significantly slower/ faster clearance). CSF from 2 pooled donors was provided from stored specimens under an observational protocol previously described [56]. As shown in Table 1, the 45 patients had an age, CD4, etc. All strains were previously serotyped to be serotype A [74].

### Strains and Media

Experiments were conducted in a genetic background of *C. neoformans* WT strain serotype A H99 (*MAT* α, ATCC 208821) and was the kind gift of J. Perfect. A complete list of strains used in this study is described in S5 Table. *Escherichia coli* DH10B (Invitrogen) was the host strain for recovery and amplification of plasmids. The fungal strains were grown in YPD medium (2% glucose, 1% yeast extract, 2% Bacto-peptone) or YPD agar medium (YPD and 2% agar). Asparagine minimum selective medium (ASN) for transformant selection and for detection of laccase production was previously described [75]. V8 juice medium were used for mating assays as described [76].

### Microarray experiments

Isolates from patients in Table 1 or the H99 lab strain were grown to mid-log phase and then transferred to asparagine medium without glucose (asparagine, 1g/L, 10 mM sodium phosphate, pH 7.4 and 0.25 g/l MgSO_4_) or human CSF pooled from 4 patients with cryptococcal meningitis and incubated for 3 h at 37°C and RNA recovered as described previously [25]. Serotype A H99-based microarrays (hybridization probe sequences described previously in [27]) with two unique probes (3 replicate features per probe) for each of 6,969 transcripts (one per locus) were used for expression profiling. Two-color co-hybridizations were performed with a common reference sample in the Cy5 channel on each array. Agilent Feature Extraction software (v. 11.5.1.1, protocol GE2_107_Sep09) estimated the median pixel intensity of each feature, which was then log2 transformed. Replicate RNA samples from CSF or starvation buffer conditions were generated in two independent experiments. The reference sample was grown in YPD and used both for loess normalization and as a comparison condition in ANOVA. Each feature signal was normalized by loess and averaged per locus (3 replicates of two probes for 1 transcript per locus). A mixed-effects ANOVA model (fixed effect of growth media, random effect of array ID) was computed, and expression difference estimates for each gene were calculated for CSF or starvation media vs. YPD. For the patient isolates (Table 1), the reference pool was created from 12 of the samples. Probe signals were summarized as the median of the 3 replicate features, then loess-normalized ratios against the cognate reference signal were averaged for the two probes per locus. A mixed-effects ANOVA model (fixed effect of survival or mortality time, random effect of Study group, two levels) was computed, and expression difference estimates for each gene were calculated for Wk2 mortality vs. survived, Wk10 mortality vs. survived, or both Wk2 and Wk10 mortality vs. survived. For both experiments, the False Discovery Rate (FDR) was estimated from raw ANOVA p-values to compensate for multiple testing of 6,969 genes. SAS and JMP/Genomics software (SAS, Cary NC) was used for statistical analysis. Data from microarray experiments were deposited on the National Library of medicine gene expression ontology (GEO) database.

### Overexpression, disruption and complementation of *ECM15*, *CAC2*, and *STL1* in *C. neoformans*

Standard methods were used for overexpression, disruption and complementation of the *ECM15*, *CAC2*, and *STL1* genes in strain H99 (*MAT*α) as described previously using two PCR-amplified fragments and a 1.3-kb PCR fragment of the URA5 gene previously described to effect a deletion within the target coding regions and was complemented using a 1-kb of up and down stream genomic fragment of the target genes [14, 77, 78]. Complementation in all cases retained the original deletion construct. Primers used in this study listed in S6 Table.

### Ecm15-mCherry and Cac2-GFP fusion proteins

The cryptococcal shuttle vector pORA-YP142 [79] was used to express a fusion between the Ecm15 protein and a synthetic mCherry protein (Cneo-mCherry), utilizing *C. neoformans* codon usage produced using standard methods [77]. The plasmid was digested with NdeI and PstI, and a PCR-amplified fragment of the H99 *ECM15* gene containing promoter region was digested with NdeI and PstI, and ligated into compatible sites to produce YP148. The plasmids were recovered, the sequences were verified, and the plasmids were linearized with SceI and transformed into *C. neoformans* H99 Matα FOA cells by electroporation using standard methods [80]. All cells preparations grown on non-selective media are assayed at the end of each experiment by simultaneous inoculation of selective and non-selective plates to verify >90% retention of plasmid. Co-localization was observed using a Leica DMI 6000B microscope with a Hamamatsu camera using LAS AF6000 ver 2.1.2 software (Leica). Predicted nuclear localization sequences were identified in the Ecm15 and Cac2 protein using cNLS-Mapper using a cut-off score of 2 [40].

### qRT-PCR experiments

*C. neoformans* strains H99 were grown on YPD or ASN without glucose media. Real-time PCR was performed using a primer sets as described in S6 Table. Reverse transcription was performed on DNase-treated RNA using the iScript kit (Bio-Rad Laboratories), according to the manufacturer’s protocol. PCRs were set up using iQ SYBR Green Supermix (Bio-Rad Laboratories), according to the manufacturer’s protocol. qRT-PCR was performed using a Bio-Rad iCycler (MyiQ2).

### Fluorescent in situ Hybridization (FISH)

Sections of a brain autopsy specimen were obtained from a 42 year old male who died of severe and diffuse *C. neoformans* infection and was reported previously [14]. FISH was performed as describe [20]. Briefly, cells were washed once with 1× PBS and fixed for 4 h with 4% w/v paraformaldehyde in PBS at 4°C. Probes were labelled at the *STL1* ORF with the C3-fluorocein (S7 Table; LGC Bioresearch). For negative control, samples were treated with RNase A (50 μg/mL) for 1 h at 37°C, prior to the hybridization step. Fixed cells were hybridized in 20 μl of hybridization buffer (0.9 M NaCl, 0.01% w/v SDS, 20 mM Tris-HCl, pH 7.2 and 20% formamide), with 5 ng of C3 fluorescein-labelled probe and incubated at 46°C for 16 h. After incubation, cells were pelleted by centrifugation and resuspended in 1 ml of prewarmed washing buffer (20 mM Tris-HCl, pH 8.0, 0.01% w/v SDS, 5 mM EDTA, 225 mM NaCl) for 30 min at 46°C. The slides were then mounted in ProLong Gold antifade reagent (Invitrogen) and observed using a Leica DMI 6000B microscope with a Hamamatsu camera using LAS AF6000 ver. 2.1.2 software (Leica). Cells were scored as positive by a blinded observer and analyzed by Fisher’s exact test.

### Measurement of capsular size and virulence studies

To induce capsule, yeast cells were grown on ASN media in a 30°C for 4 days. Capsule was measured by microscopy after the fungal cells were suspended in India ink [81], and laccase by melanin production on nor-epinephrine agar. Virulence studies were conducted according to a previously described intravenous mouse meningoencephalitis model [82] using 10 CBA/J mice for each *C. neoformans* strain. All experimental procedures were conducted under a protocol approved by the Institutional Animal Care and Use Committee of the Intramural Research Program of the NIAID, NIH.

### Chromatin Immunoprecipitation (ChIP) assay

The ChIP assay was adapted and modified from a previously described protocol [83] for *C. neoformans*. PCR detection of the *STL1* was performed using a primer sets as described in S5 Table. ‘Input’ DNA was used as a positive control, consisting of unprecipitated genomic DNA as a loading control and to show intact function of each primer set and PCR reaction.

### Promoter deletion studies

5’-truncated promoter sequences were obtained by PCR using a single reverse primer and different forward primers (−1000, −500, −20 bp from the transcript start site) carrying BglII and NdeI restriction sites (S5 Table). The amplified fragments were inserted in the upstream of the GFP gene coding region of the plasmid, producing a series of STL1pro-STL1-GFP vectors. After verification by sequence analysis, the confirmed constructs or linear amplified fragments containing the indicated upstream region, *STL1* reading frame and GFP marker and terminator as indicated were transformed into WT or *cac2*Δ strains of *C. neoformans*. Integrated constructs were confirmed by uncut Southern blots [83]. These strains subjected to microscopy (DIC, GFP fluorescence) or flow cytometry (FITC) to determine the promoter activity of *STL1* under glucose or starvation conditions.

### Statistics

Errors were expressed as standard error of the mean (SEM). Fluorescent positive cells (Fig 1) were scored in a blinded fashion and analyzed by Fisher’s exact test.

Calculations: Statistics available from the ANOVA output (JMP/Genomics version 8.0) include mean square error (MSE), error degrees of freedom (DDF), model degrees of freedom (NDF) and the proportion of the variance accounted for by the model (R^2^). From these we derive:

SSE (sum of squares error) = MSE * DDF

SSM (sum of squares model) = SSE * R^2^ / (1 - R^2^)

MSM (mean square model) = SSM / NDF

F-ratio = MSM/MSE

Raw p-values are retrieved from the cumulative probability distribution of the F-ratio using the parameters F-ratio, NDF and DDF (e.g., with the SAS function PROBF)

N (total samples) = NDF + DDF + 1

C (number of groups) = NDF + 1

A hypothetical increase of n samples per group by iteration i (steps) leads to an increase in DDF

DDF(ni) = N + 2*n*i – C

which propagates to recalculations of SSE, SSM, F-ratio and p-value for i=1,2,3… iterations.

## Acknowledgements

We acknowledge all members of JMP Technical Support team, SAS Institute, for assistance with power calculation formulas. We thank J. Powell (Bioinformatics and Molecular Analysis Section, CIT, NIH) for assistance with microarray data management and informatics.

## Supporting Information

**S1 Fig. Metabolic reprogramming of carbohydrate metabolism in *C. neoformans* in response to incubation in fungal infected human CSF or starvation.** Pathway schemes demonstrate differential expression ratios of *C. neoformans* after incubation in human fungal infected CSF (CSF) or in starvation conditions (starv) relative to mid log cells grown in YPD of genes expressing enzymes active in b-oxidation, glyoxylate cycle or gluconeogenesis. Adapted from Derengowski et al.

**S2 Fig. Expression profiling demonstrates transcriptional expansion of genes related to transporter activity (GO function).** Transcriptional profiling of *C. neoformans* cells as in Fig 1A displayed by most common GO function terms. Percent of annotated GO terms relative to total count of gene annotations (left upper panel), starvation (right upper panel), CSF alone (right lower panel) or both CSF and starvation (right lower panel).

**S3 Fig. Prediction of putative nuclear localization signals.** Ecm15 (A) and Cac2 (B) putative nuclear localization signals were determined by cNLS Mapper.

**S4 Fig. Spotting assay for growth.** Indicated strains were diluted to an A_600_ of 1.0, and 1:5 serial dilutions (5μl) were plated on ASN media containing the indicated substrates (0.03%) and incubated at 30°C for 5 to 10 days.

**S5 Fig. Chromatin immunoprecipitation (ChIP) survey of Cac2-promoter complexes.** Chromatin immunoprecipitation (ChIP) of a Cac2-promoter complex. Nuclear extract from induced cryptococcal cells was immunoprecipitated as described in Methods and assayed by PCR for the presence of the indicated regions of *STL1* or *FRE7* promoter sequences or control promoter sequences of *ACT1*.

**S6 Fig. Effects of serial deletion of the *STL1* 5′-promoter (−1000 from the transcript start site) on basal transcriptional activity in *C. neoformans*.** A plasmid containing *STL1* ORF with indicated regions of the *STL1* 5′-promoter GFP fusion gene were transformed into WT or *cac2*Δ strains of *C. neoformans* and grown to mid-log in YPD (Glu +) and ASN salts without glucose (Glu -) media and the population subjected to microscopy (DIC, GFP fluorescence) or flow cytometry (FITC). Empty vector used as a control. Bar=5μm.

**S7 Fig. qRT-PCR of *STL1*.** qRT-PCR of STL1 of Y813 strain grown to mid-log in YPD (Glu +) and ASN without glucose (Glu -) media. N = 3 independent experiments, Student’s t-test, *; p < 0.05.

**S8 Fig. Mating assay**. Indicated MATα strains were co-incubated with a MATa mating partner (strain KN99) on nitrogen limiting mating media (V8 agar) for 1 week at 25°C. The edges of the mating mixtures were photographed (40x).

**S9 Fig. Capsule thickness across mutant strains in the *C. neoformans*.** Indicated strains were incubated on YPD (Glu +) and ASN without glucose (Glu -) agar for 3 days at 30°C and examined by India ink microscopy. Bar = 5 μm.

**S10 Fig.** qRT-PCR of *STL1* of indicated strains grown to mid-log in YPD (Glu +) and ASN media containing 0.03 % acetate (without glucose). N = 3 independent experiments, Student’s t-test, *; p < 0.05, ***; p<0.001.

**S1 Table.** Genome-wide expression level changes after transition from YPD to CSF

**S2 Table.** Genome-wide expression level changes after transition from YPD to starvation media

**S3 Table.** Genome-wide expression changes between isolates of patients who died versus survived at 10 weeks.

**S4 Table.** List of genes suppressed by both *ECM15* and *CAC2* determined by genome-wide expression analysis

**S5 Table.** Strains used in this study.

**S6 Table.** Primers used in this study.

**S7 Table.** FISH probes used in this study.

